# T cell-specific loss of IRF1 results in defective CD8 T cell activation and antitumor immunity

**DOI:** 10.64898/2026.05.14.725206

**Authors:** Lulu Shao, Hridesh Bannerjee, Ecem Unal, Isha Mehta, Jishnu Das, A. Rouf Banday, Larry Kane, Saumendra N. Sarkar

## Abstract

Interferon regulatory factor 1 (IRF1) has long been recognized as a tumor suppressor; however, recent studies have revealed context-specific and sometimes opposing roles in cancer progression. Here, we describe a T cell–specific mechanism underlying the antitumor activity of IRF1. Unlike germline *Irf1*-deficient mice, T cell–specific loss of IRF1 does not lead to a deficiency in cytotoxic CD8⁺ T cells. Nevertheless, tumor burden remains elevated in these mice, associated with reduced CD8⁺ T cell infiltration driven by impaired activation and proliferation in the absence of IRF1. Transcriptomic analysis of activated *Irf1*-deficient T cells identified NFATc1 as a key gene significantly downregulated upon IRF1 loss. Analysis of human melanoma datasets further corroborated this finding, highlighting a previously unappreciated role for IRF1 in regulating T cell activation and antitumor immunity.

## INTRODUCTION

Interferon regulatory factors (IRFs) were discovered and characterized as transcriptional regulators of type I interferons (IFNs) and IFN-inducible genes. Specific members of the IRF family are involved in the induction of IFN, lymphocyte development and oncogenic signaling (1–3). IRF1 has been known as an antitumor gene based on human leukemia (4–6) as well as other studies including *Irf1*-deficient mouse studies (7,8). However, as we have shown recently the antitumor function of IRF1 is highly context-specific. As opposed to its antitumor function in the immune cells, IRF1 plays a pro-tumorigenic function in the tumor cell. In the tumor cells IRF1 is required for transcriptional upregulation of PD-L1 to block antitumor T cell response and evade host immune surveillance (9) and preferential inhibition of IRF1 in the tumor cells led to significantly reduced tumor growth (10).

Additionally, *Irf1^-/-^* mice are known to exhibit significant deficiency in CD8^+^ T cells (11). It has been proposed that IRF-1 regulates gene expression in thymic stromal cells, and in developing thymocytes, it is required for lineage commitment and selection of CD8^+^ thymocytes. As a result, the germline *Irf1*-deficient mice are to some extent immunodeficient due to the lack of CD8^+^ T cells (11,12). Therefore, to characterize the immune cell specific roles of IRF1 in mediating antitumor immunity we created various tissue-specific IRF1-deficient mice and characterized tumor growth. We found that unlike the germline IRF1-deficient mice, the naive CD8^+^ T cell population is restored in the T cell-specific IRF1-deficient mice. However, CD8^+^ T cell activation and tumor infiltration are impaired in the absence of IRF1 in T cells. Transcriptomic analysis identified reduced expression of NFAT family transcription factors in CD8^+^ T cells. This finding is consistent in the context of human tumor samples, where higher expression of IRF1 was correlated with increased expression of NFATs. This study, therefore, establishes a critical role of IRF1 in T cell receptor (TCR) signaling in CD8^+^ T cells in the context of antitumor immune response.

## MATERIALS AND METHODS

### Cell line

B16-F10 cells were purchased from ATCC, and cells were used within 10 generations. Cells were maintained in RPMI-1640 medium (Corning) supplemented with 10% FBS (R&D systems) and 100 U/mL penicillin/streptomycin (Gibco).

### Generation tissue-specific *Irf1*-KO mice and tumor model

Mice used in this study were housed in specific pathogen-free conditions with the approval of the University of Pittsburgh Institutional Animal Care and Use Committee (IACUC, protocol number: IS00022132). *Irf1^fl/fl^* mice were a kind gift from Dr. Michael Diamond (Washington University). *EIIa^Cre^* (B6.FVB-Tg(EIIa-Cre)C5379Lmgd/J, cat# 003724), *Cd4^Cre^* (B6.Cg-Tg(Cd4-Cre)1Cwi/B fluJ, cat# 022071) and *Lyz2^Cre^* (B6.129P2-Lyz2tm1(Cre)Ifo/J, cat# 004781) mice were purchased from the Jackson laboratory. The *Irf1^fl/fl^* mice were bred with *EIIa^Cre^*, *Cd4^Cre^*and *Lyz2^Cre^* mice respectively to generate full-body, T cell- and myeloid cell-specific *Irf1*-KO mice. PCR genotyping was regularly conducted to confirm and maintain all the mouse lines, and the primers used in the study were listed in supplemental Table 1. The removal of IRF1 was tested in splenocytes and isolated T cells from those mice by immunoblotting and flow cytometry with IRF1 antibody (CST, cat# 8478P) and IRF1-PE antibody (CST, cat# 12732), respectively. For B16-F10 tumor model, mice were used at 8-10 weeks old and intradermally injected with 5 × 10^5^ B16-F10 cells. Tumor volumes were measured and calculated 3 times a week with the formula V = 0.52 × width^2^ × length.

### Immuno-profile of thymus, spleen and lymph nodes of tissue-specific *Irf1*-KO mice

Naïve *Irf1^fl/f^*, *EIIa^Cre^Irf1^fl/fl^*, *Cd4^Cre^Irf1^fl/fl^*, *Lyz2^Cre^Irf1^fl/fl^*mice were euthanized at 10-week-old, 2 males and 2 females for each group. Thymus, spleen, axillary lymph node (LN), brachial LN and inguinal LN were collected from those mice, then mechanically smashed between two frosted glass slides and filtered through 70 µM cell strainer to get single-cell suspension. Cells were resuspended in FACS buffer and stained with the following fluorescence-conjugated Abs: anti-CD3 (clone 145-2C11, BioLegend), anti-CD4 (clone RM4-5, eBioscience), anti-CD8 (clone 53-6.7, BioLegend), anti-F4/80 (clone BM8, BioLegend), anti-CD11b (clone M1/70, BioLegend), anti-CD11c (clone N418, BioLegend), anti-NK1.1 (clone PK136, BioLegend), anti-I-A/I-E (clone M5/114.15.2, BioLegend), anti-CD19 (clone 6D5, BioLegend), anti-CD25 (clone PC61.5, eBioscience), anti-Foxp3 (clone FJK-16s, eBioscience). Cells stained for Foxp3 were fixed and permeabilized with the Foxp3/Transcription Factor Staining Buffer Set (eBioscience). BD LSRFortessa cytometer and FACSDiva software (BD Bioscience) were used to collect the data. Color compensation and analyses were performed using FlowJo software (FlowJo). We designated T regulatory cells (Treg) = CD3^+^, CD4^+^, CD25^+^, Foxp3^+^.

### Tumor infiltrating lymphocytes analysis

Tumors were collected on day 12 post-injection, and injected with 0.5 – 1 mL of tumor digestion mixture [2 mg/mL collagenase IV (Thermo Fisher Scientific), 200 μg/mL Hyaluronidase V (Sigma-Aldrich), and 4 U/mL DNase I (Sigma-Aldrich)], then incubated at 37°C for 20 minutes. Digested tumors were dissociated between two frosted glass slides and filtered through 70 µM cell strainer. Tumors cells were subjected to flow staining with the following fluorescence-conjugated Abs: anti-CD3, anti-CD4, anti-CD8, anti-CD25, anti-CD45 (clone 30-F11, BioLegend), anti-Tim3 (clone RMT3-23, BioLegend), anti-Lag3 (clone C9B7W, BioLegend), anti-PD-1 (clone 29F.1A12, BioLegend), anti-Foxp3, anti-Ly6c (clone HK1.4, eBioscience), anti-Ly6G (Gr-1) (clone RB6-8C5, eBioscience). We designated exhausted T cells (ExT) = CD3^+^, CD8^+^, PD-1^high^, Tim3^+^, Lag3^+^; T regulatory cells (Treg) = CD3^+^, CD4^+^, CD25^+^, Foxp3^+^; granulocytic myeloid derived suppressor cells (GrMDSC) = CD45^+^, CD11b^+^, Gr-1^+^ and monocytic MDSC = CD45^+^, CD11b^+^, Ly6c^+^.

### Examination of T cell activation and proliferation

Naïve CD8^+^ T cells were isolated from splenocytes of *Irf1^fl/fl^*and *Irf1^fl/fl^Cd4^Cre^* mice using MojoSort^TM^ mouse CD8 naïve T cell isolation kit (BioLegend) followed the manufacturer’s instructions. The isolated T cells were activated with plate bound CD3 Ab T cell activation method. 100 µL of sterile PBS containing 3 µg/mL of purified CD3 Ab (BioLegend) was added into each well of a flat-bottom 96-well cell culture plate and incubated at 37°C overnight. The isolated naïve CD8^+^ T cells were resuspended with R10 medium containing 2 µg/mL of purified CD28 Ab (clone 37.51, BioLegend) and 50 U/mL mouse IL-2 (Pepro Tech Inc.), and added into each well after the removal of CD3 Ab-containing PBS, then incubated at 37°C with 5% CO_2_. Cells were harvested at different time points and subjected to flow staining with anti-CD25, anti-CD69 (clone H1.2F3, BioLegend), and anti-Ki67 (clong B56, BD Horizon). Foxp3/Transcription Factor Staining Buffer Set was used for Ki67 staining.

To examine cytokine production in CD8^+^ T cells, cells were activated either by the method mentioned above, or by 100 ng/mL phorbol 12-myristate 13-acetate (PMA) and 500 ng/mL ionomycin (Thermo Fisher Scientific) overnight, then incubated with BD GolgiPlug protein transport inhibitor (1:1000) (BD Biosciences) for 4 hours. After incubation, cells were harvested for intracellular cytokine staining with anti-IFNγ (clone XMG1.2, BioLegend) and anti-TNFα (clone MP6-XT22, BioLegend) as described above.

### Differential Gene Expression Analysis

Naïve CD8^+^ T cells were isolated from splenocytes of *Irf1^fl/fl^*and *Irf1^fl/fl^Cd4^Cre^* mice and activated by anti-CD3 and anti-CD28 as described above. Cells were harvested at 0, 6, 12 and 24 hours, frozen by dry ice, and subjected to RNA-sequencing by Novogene.

Raw sequencing reads were subjected to quality control and adapter trimming. Transcript-level quantification was performed using kallisto (13), with the mm10 (UCSC) mouse reference transcriptome. Transcript abundance estimates were imported into sleuth (14) for differential expression analysis, incorporating quantification uncertainty. Differentially expressed transcripts were identified by comparing Irf1-deficient CD8⁺ T cells to control cells at each stimulation time point (6 h, 12 h, and 24 h). Transcripts were considered significantly differentially expressed using a Benjamini–Hochberg adjusted q-value threshold of < 0.2, consistent with exploratory transcriptomic analyses.

Over-representation analysis (ORA) and Gene Set Enrichment Analysis (GSEA) (15) were performed using the clusterProfiler (16) package to identify enriched biological pathways. Pathway annotations were derived from the KEGG (17) and Reactome (18) databases. Heatmaps and principal component analysis (PCA) were generated using normalized transcript counts to assess global transcriptional changes and sample clustering across genotypes and stimulation time points.

### Detection the expression of NFATc1 in CD8^+^ T cells

Naïve CD8^+^ T cells were isolated from splenocytes of *Irf1^fl/fl^*and *Irf1^fl/fl^Cd4^Cre^* mice and activated by anti-CD3, anti-CD28 and mouse IL-2 as described above. Cells were harvested at 0, 0.5, 2 and 6 hours post-treatment, subject to flow staining with anti-NFATc1 (clone 7A6, BioLegend) using Foxp3/Transcription Factor Staining Buffer Set.

### Single-cell RNA-seq analysis of human melanoma T cells

Single-cell RNA-seq data from a human melanoma cohort (GEO accession: GSE148190) were analyzed using Seurat (v4) (19). Raw count matrices were processed using standard quality-control procedures, including filtering of low-quality cells and normalization via log-transformation of library-size–scaled counts. Highly variable genes were identified, and dimensionality reduction was performed using principal component analysis followed by Uniform Manifold Approximation and Projection (UMAP) (20).

T cells were identified based on canonical lineage markers (e.g., CD3D, CD3E, CD2) and subsetted for downstream analyses. UMAP embeddings were used to visualize overall cellular composition and to assess the distribution of IRF1 and NFATC1 expression within the T-cell compartment.

To quantify the association between IRF1 and NFATC1 while accounting for single-cell sparsity and inter-patient variability, we implemented a per-sample pseudobulk strategy in which each patient sample served as the unit of analysis. For each sample, mean log-normalized gene expression was calculated across T cells. Pseudobulk summarization was performed for (i) all T cells, (ii) CD8⁺ T cells (defined by CD8A and/or CD8B expression), and (iii) CD4⁺ T cells (defined by CD4 expression). Sample-level mean IRF1 and NFATC1 expression values were visualized using scatter plots and assessed for association using Spearman rank correlation with two-sided p-values.

To determine whether this relationship extended across the NFAT transcription factor family, Spearman correlations between IRF1 and NFATC1–NFATC5 were computed using the same pseudobulk T-cell samples. Correlation coefficients and nominal p-values are reported.

Finally, to evaluate whether NFATC1 expression was associated with differences in T-cell state composition, samples were stratified into NFATC1-High and NFATC1-Low groups based on the median pseudobulk NFATC1 expression across all T cells within the cohort. Downstream comparisons of T-cell state distributions were performed between these groups.

### Statistical analysis

Animal experiment results shown were pooled samples from at least two repeated studies. The *in vitro* studies were repeated at least three times. For each data point mean and SEM were plotted, statistical significance calculated either by unpaired Student’s t-test or two-way ANOVA with Sidak’s multiple comparison test as appropriate. The survival curves are compared statistically using the log rank test and Gehan-Breslow-Wilcoxon test. The results represented as * P < 0.05, ** P < 0.01, *** P < 0.001, **** P < 0.0001.

## RESULTS

### T cell-specific loss of IRF1 does not affect T cell development

Given the immunocompromised state of germline IRF1-deficient mice (*Irf1^−/−^*) due to the loss of CD8 T cells, we generated and immunoprofiled various tissue specific IRF1-deficient mice. *Irf1^fl/fl^* mice were bred with *EIIa^Cre^*, *Cd4^Cre^* and *Lyz2^Cre^* mice to generate germline IRF1-deficient (*Irf1^fl/fl^EIIa^Cre^*), T cell-specific IRF1-deficient (*Irf1^fl/fl^Cd4^Cre^*), and myeloid cell-specific IRF1-deficient (*Irf1^fl/fl^Lyz2^Cre^*) mice. Splenocytes and isolated T cells from these mice were treated with mouse IFNγ to confirm the deletion of IRF1 by western blotting (Fig. S1A). IRF1 expression levels were also compared by intracellular flow cytometry in CD11b^+^ and CD3^+^ cells among tissue-specific *Irf1*-deficient mice with *Irf1^fl/fl^* as controls. As shown in Fig. S1B, CD3^+^ T cells from *Irf1^fl/fl^* and *Irf1^fl/fl^Lyz2^Cre^*mice had comparable levels of IRF1 expression, whereas *Irf1^fl/fl^EIIa^Cre^*and *Irf1^fl/fl^Cd4^Cre^* mice showed minimal IRF1 expression in CD3^+^ T cells. CD11b^+^ myeloid cells from *Irf1^fl/fl^Lyz2^Cre^*mice showed substantially reduced IRF1 expression compared to cells from *Irf1^fl/fl^*mice, whereas IRF1 expression was also reduced to some extent in CD11b^+^ myeloid cells from *Irf1^fl/fll^Cd4^Cre^* mice (Fig. S1B). These data demonstrated the successful deletion of IRF1 protein expression in T cells of germline and T cell-specific *Irf1*-deficient mice.

Next, we compared the immune cell composition of the thymus, spleen and lymph nodes in *Irf1^fl/fl^* and tissue-specific *Irf1*-deficient mice. As expected, we found that *Irf1^fl/fl^EIIa^Cre^* mice had significantly less CD8^+^ T cells in thymus, spleen and lymph nodes but slightly more CD4^+^ T cells in spleen and lymph nodes compared to other 3 groups of mice (red bars in Fig. 1A-B). However, T cell-specific *Irf1-*deficient mice presented minimal differences in various immune cell populations compared to *Irf1^fl/fl^* mice (green bars in Fig. 1C and Fig. S1C-S1E). The *Irf1^fl/fl^EIIa^Cre^* mice also showed significantly less NK cells and more Tregs in spleen compared to other groups of mice (Fig. 1C and Fig. S1F). However, the higher percentage of Tregs in CD3^+^ T cells was related to the enrichment of CD4^+^ T cells in total population of T cells, as the percentage of Tregs did not show any significant changes in CD4^+^ T cell composition (Fig. S1E-S1G).

**Fig. 1:**
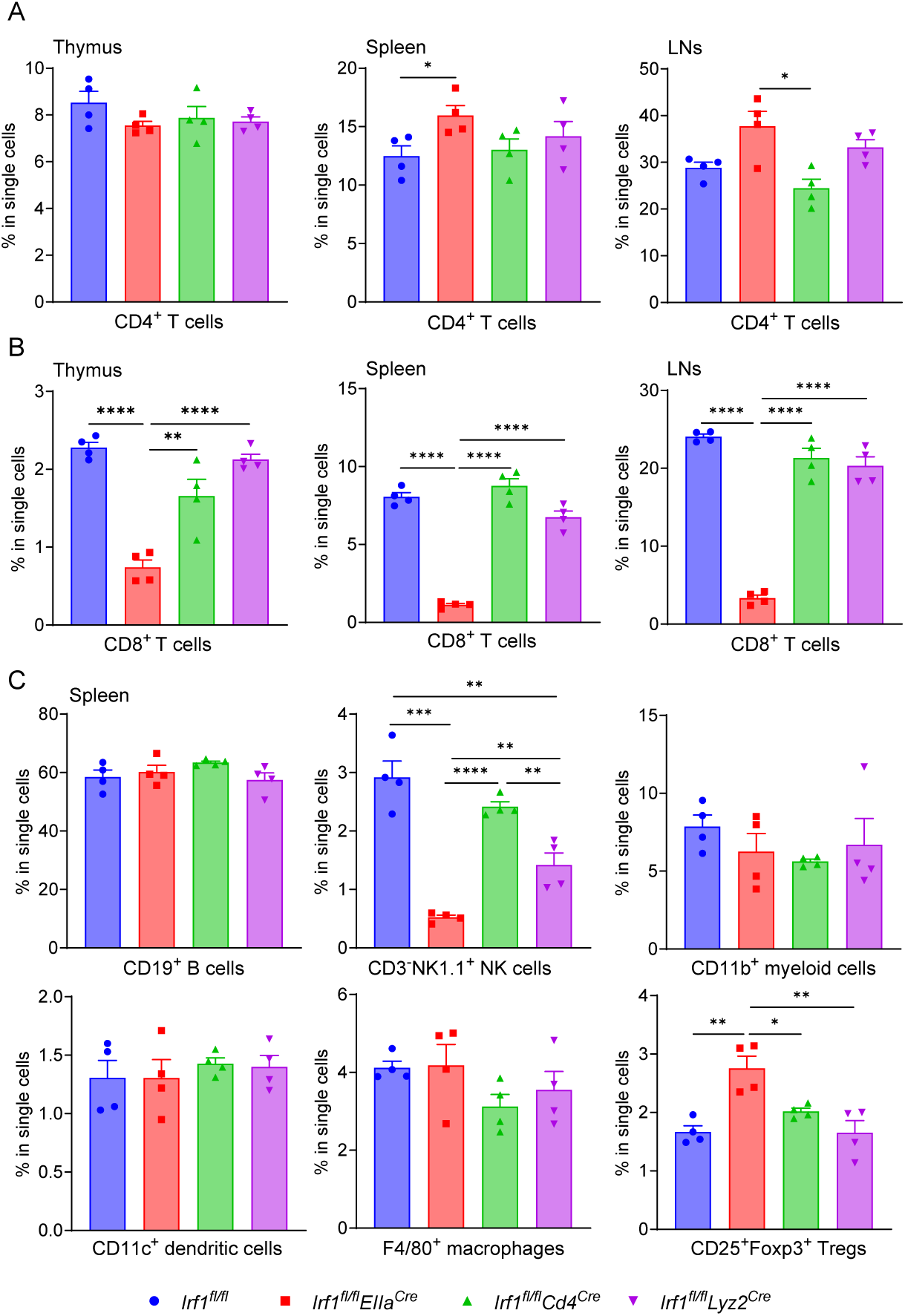
Loss of IRF1 in T cells does not affect the development of T cells. (A) Percentages of CD4^+^ T cells and (B) CD8^+^ T cells in thymus, spleen, and lymph nodes (LNs). The thymus, spleen and lymph nodes were collected from 9-10 weeks old of *Irf1^fl/fl^*, *Irf1^fl/fl^EIIa^Cre^*, *Irf1^fl/fl^Cd4^Cre^*, and *Irf1^fl/fl^Lyz2^Cre^* mice, 2 females and 2 males in each group (n=4). Single cell suspensions were stained with anti-CD3, anti-CD4, anti-CD8, anti-CD25, and anti-Foxp3 Abs. Data was collected by BD LSRFortessa cytometer and analyzed by FlowJo. (C) Percentages of B cells, NK cells, myeloid cells, dendritic cells, macrophages and T regulatory (Tregs) in spleen. The spleen was collected from 9-10 weeks old of *Irf1^fl/fl^*, *Irf1^fl/fl^EIIa^Cre^*, *Irf1^fl/fl^Cd4^Cre^*, and *Irf1^fl/fl^Lyz2^Cre^*mice, 2 females and 2 males in each group (n=4). Single cell suspensions were stained with anti-F4/80, anti-CD11b, anti-CD11c, anti-NK1.1, anti-I-A/I-E, anti-CD19 Abs. Data was collected by BD LSRFortessa cytometer and analyzed by FlowJo. Mean and SEM were plotted for each group of mice, and statistical significance was calculated by multiple unpaired t-tests and represented as * P < 0.05, ** P < 0.01, *** P < 0.001, **** P < 0.0001.

### IRF1 in T cells is important to suppress tumor progression

Given the established tumor suppressive function of IRF1 and our previous findings that tumor cell intrinsic role of IRF1 to evade antitumor immune response by upregulating PD-L1 expression, we sought to examine the T cell and myeloid cell specific function of IRF1 in antitumor immunity using a syngeneic mouse melanoma model. B16-F10 cells were intradermally injected into the back of all four groups of mice, and tumor size and survival were monitored. Consistent with previous studies, full-body *Irf1*-deficient mice (*Irf1^fl/fl^EIIa^Cre^*) showed significantly accelerated tumor growth compared to the *Irf1^fl/fl^* group of mice. Surprisingly, T cell-specific *Irf1*-deficient (*Irf1^fl/fl^CD4^Cre^*) mice also showed significantly accelerated tumor growth compared to control group with (Fig. 2A), while myeloid cell-specific *Irf1*-deficient (*Irf1^fl/fl^Lyz2^Cre^*) mice had similar tumor growth patterns as the *Irf1^fl/fl^* group (Fig. 2A). According to the survival curves, *Irf1^fl/fl^EIIa^Cre^* and *Irf1^fl/fl^CD4^Cre^* mice exhibited significantly reduced survival than the other two groups (Fig. 2B). Given that unlike germline *Irf1*-deficient mice, T cell-specific *Irf1*-deficient mice showed no defect in various immune cell populations (Fig. 1), these results indicated that IRF1 expression in T cells is important for effective antitumor responses of the host.

**Fig. 2:**
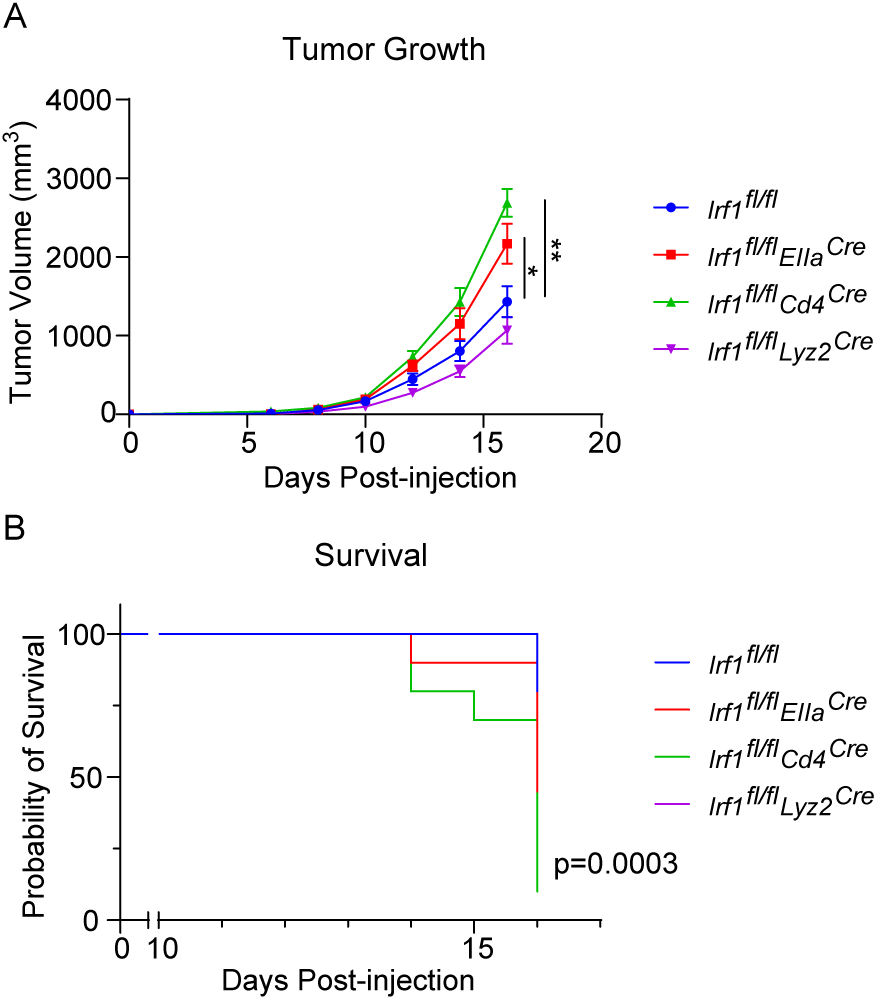
T cell specific loss of IRF1 impairs antitumor immunity. (A) The tumor growth and (B) survival B16-F10 tumor-bearing tissue-specific *Irf1*-KO mice. 5 × 10^5^ of B16-F10 cells were intradermally injected into 4 groups of mice, followed by tumor growth measurements. Tumor volumes were measured and calculated 3 times a week with the formula V = 0.52 × width^2^ × length. Combined data from two biological repeats. *Irf1^fl/fl^* mice, n=10; *Irf1^fl/fl^EIIa^cre^*mice, n=8; *Irf1^fl/fl^Cd4^cre^* mice, n=10; *Irf1^fl/fl^Lyz2^cre^*mice, n=10. The growth curves represent corresponding mean values with SEM. Multiple unpaired t-tests were used to calculate the P value for growth curves. The survival curves are compared using the log rank test and Gehan-Breslow-Wilcoxon test. The results were represented at * P < 0.05, ** P < 0.01.

### Tumors from *Irf1^fl/fl^CD4^Cre^* mice have reduced CD8^+^ T cell infiltration

To investigate the T cell specific role of IRF1, we collected B16-F10 tumors from *Irf1^fl/fl^* and all tissue-specific *Irf1*-deficient mice on post-injection day 12 and analyzed the profiles of tumor-infiltrating immune cells, especially T cells, using flow cytometry. Tumors from all 4 groups of mice had comparable amounts of infiltrated CD3^+^ T cells (Fig. 3A). However, all 3 groups of tissue-specific *Irf1*-deficient mice tumors had significantly less infiltrating CD8^+^ T cells (Fig. 3B). Tumors from full-body *Irf1*-deficient mice had very little CD8^+^ T cells and limited percentages of CD4^+^ T cells (Fig. 3B-3D and Fig. S2A). Although the infiltrating CD8^+^ T cells from T cell-specific *Irf1*-deficient mice tumors were statistically fewer compared to *Irf1^fl/fl^* group, the proportions in the CD3^+^ population were substantially higher than the full-body *Irf1*-deficient group (Fig. 3B) and significantly higher in the frequency of total cells (Fig. S2A). We also noticed that there were significant differences in CD8^+^ T cells percentages between *Irf1^fl/fl^* and myeloid cell-specific *Irf1^fl/fl^Lyz2^Cre^*groups (Fig. 3B and Fig. S2A), but similar levels of MDSCs (Fig. S2C).

**Fig. 3:**
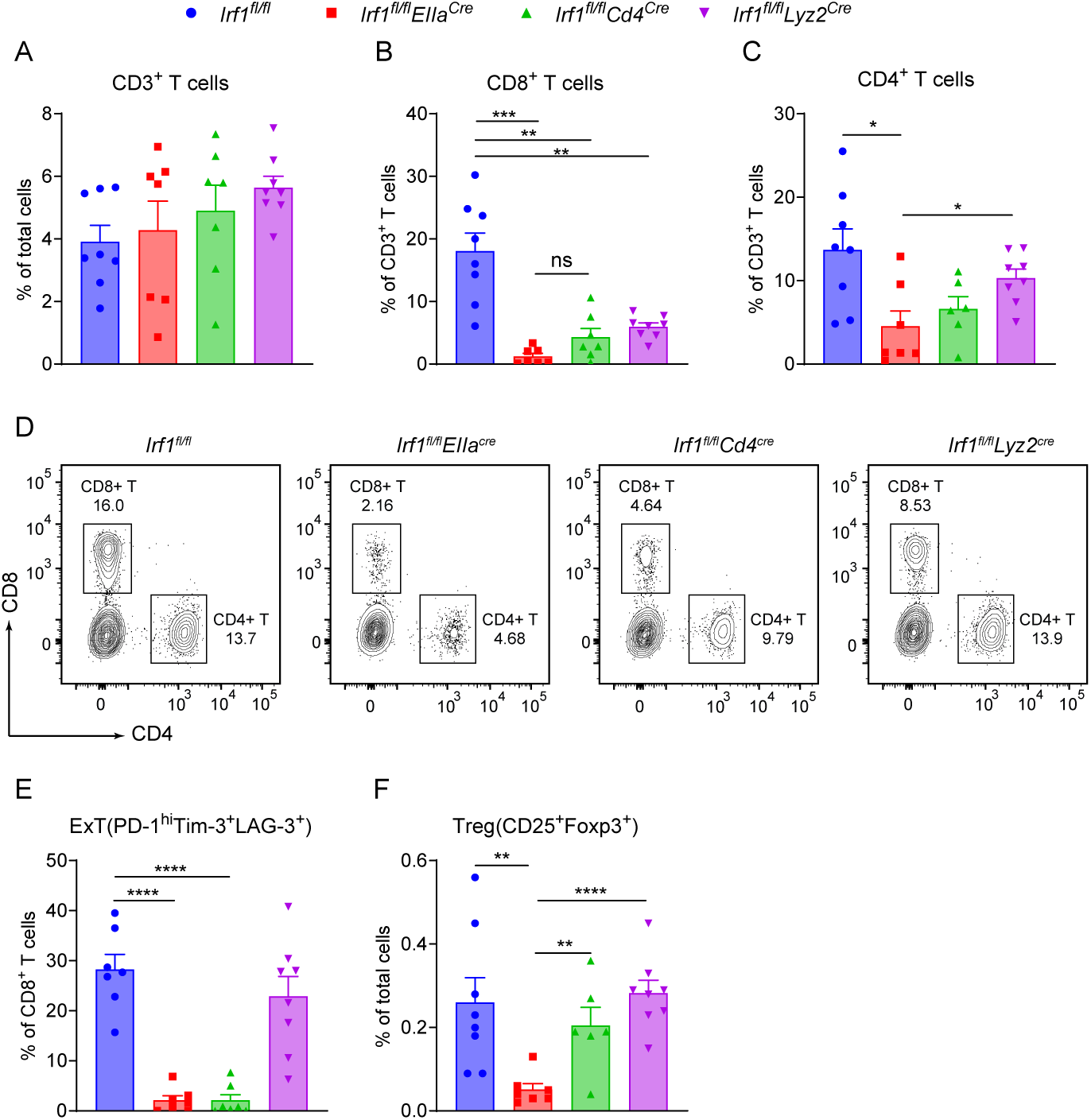
Loss of IRF1 in T cells decreases CTL infiltration in tumors. B16-F10 cells were intradermally injected into different tissue-specific *Irf1*-KO mice, and tumors were collected on post-injection day 12. Tumors were subjected to enzyme digestion and dissociated into single-cell suspensions, then stained with anti-CD3, anti-CD4, anti-CD8, anti-CD25, anti-Tim3, anti-Lag3, anti-PD-1, anti-Foxp3 Abs. Data was collected by flow cytometry and analyzed by FlowJo. (A) Percentage of CD3^+^ T cells in tumor single-cell suspension. Percentages of (B) CD8^+^ T cells and (C) CD4^+^ T cells in tumor infiltrated CD3^+^ T cells. (D) Flow cytogram of CD4^+^ and CD8^+^ T cells in tumor infiltrated CD3^+^ cell population. (E) Percentage of exhausted T cells in tumor infiltrated CD8^+^ T cells. (F) Percentage of Treg in tumor single-cell suspension. Combined data from two biological repeats. *Irf1^fl/fl^* mice, n=8; *Irf1^fl/fl^EIIa^cre^*mice, n=7; *Irf1^fl/fl^Cd4^cre^* mice, n=7; *Irf1^fl/fl^Lyz2^cre^*mice, n=8. Multiple unpaired Student’s t tests were used and the results were represented as * P < 0.05, ** P < 0.01, *** P < 0.001, **** P < 0.0001.

Tumors from T cell-specific *Irf1^fl/fl^CD4^Cre^*mice exhibited increased CD8^+^ T cell infiltration relative to full-body *Irf1*-deficient (*Irf1^fl/fl^EIIa^Cre^*) tumors, but the percentage of ExT cells among CD8^+^ T cells was nearly identical between the two groups (Fig. S2A and Fig. 3E). In addition, there were significantly higher numbers of infiltrated Tregs in T cell-specific *Irf1^fl/fl^CD4^Cre^* tumors compared to *Irf1^fl/fl^EIIa^Cre^* tumors (Fig. 3F and Fig. S2A-S2B), which agreed with more tumor infiltrated CD4^+^ cells in the first group. Taken together, these data suggested that loss of IRF1 in T cells primarily affects tumor infiltration of CD8^+^ T cells rather than CD4^+^ T cells during antitumor responses.

### IRF1 is needed for CD8^+^ T cell activation and proliferation

The reduced CD8^+^ T cell infiltration observed in tumors from *Irf1^fl/fl^CD4^Cre^* mice may be caused by deficient CD8^+^ T cell activation or infiltration. To examine whether loss of IRF1 in T cells would affect CD8^+^ T cell activation, naïve CD8^+^ T cells were isolated from splenocytes of *Irf1^fl/fl^*and T cell-specific *Irf1*-KO mice, and stimulated with CD3, CD28 Ab and mouse IL-2. The expression levels of T cell activation markers, CD25 and CD69, were evaluated at different time points. We found that the percentages of CD25^+^CD69^+^ CD8^+^ T cells were higher in wild type T cells compared to *Irf1*-KO T cells at all 4 time points (Fig. 4A and Fig. S3A) and showed statistically significant differences at 12 hours post-stimulation (Fig. 4B). The expression levels of Ki67 in CD8^+^ T cells exhibited similar pattern, and there was significantly higher frequency of Ki67 ^+^CD8^+^ T cells in *Irf1^fl/fl^* group than *Irf1^fl/fl^CD4^Cre^* group at 48 hours post-treatment (Fig. 4C-4D). In addition, intracellular expression levels of two essential antitumor cytokines, IFN- γ and TNF- α, were assessed in these two groups of T cells. We found that loss of IRF1 impaired IFN-γ expression, but not TNF-α production in the CD8^+^ T cells (Fig. 4E). This was only observed in CD3, CD28 Ab and mouse IL-2 treated, not PMA/ionomycin treated T cells (Fig. S3B). As CD3/CD28/IL-2 activation mimics physiological TCR signaling, while PMA/ionomycin is a pharmacological shortcut that bypasses the TCR altogether (21), our results indicate that the loss of IRF1 in T cells impairs downstream TCR signaling in CD8^+^ T cells.

**Fig. 4:**
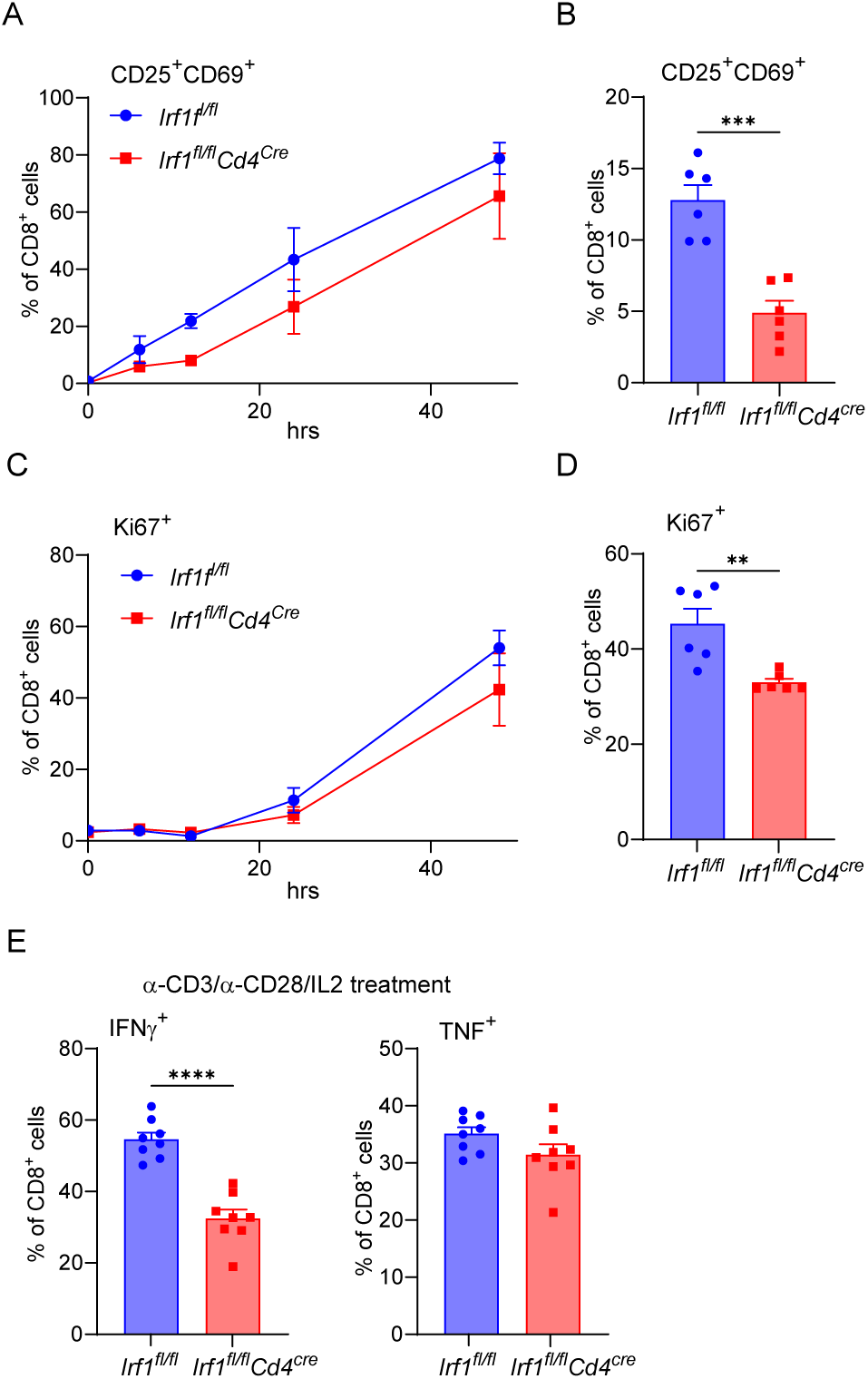
CD8^+^ T cell activation and proliferation are affected by the absence of IRF1. Naïve CD8+ T cells were isolated from splenocytes of *Irf1^fl/fl^* and *Irf1^fl/fl^Cd4^Cre^*mice using MojoSort mouse CD8 naïve T cell isolation kit, then treated with mouse anti-CD3 Ab, anti-CD28 Ab and IL-2, and cells were harvested and subjected to flow staining at 6, 12, 24, and 48 hours. (A) The expression of activation markers, CD25 and CD69, were measured at 6, 12, 24 and 48 hours after treatment by flow cytometry. (B) Percentage of CD25^+^CD69^+^ in CD8+ T cells at 12 hours post-treatment. (C) The expression of proliferation marker, Ki67, was measured at 6, 12, 24 and 48 hours after treatment by flow cytometry. (D) Percentage of Ki67^+^ in CD8^+^ T cells at 48 hours post-treatment. (E) Percentages of IFNγ^+^ or TNFα^+^ in CD8^+^ T cells at 48 hours post-treatment. The experiments were biologically repeated 3 times. For each data point, mean and SEM were plotted, and statistical significance was calculated by unpaired t-test, and presented as ** P < 0.01, *** P < 0.001, **** P < 0.0001.

### Transcriptomic analysis of IRF1-deficient CD8^+^ T cells shows defective TCR and NFAT Signaling

The TCR signaling pathway is a complex cascade, which involves hundreds of genes (22). To define the molecular mechanisms underlying impaired CD8⁺ T cell activation in the absence of IRF1, we performed transcriptomic analysis. Naïve CD8^+^ T cells isolated from splenocytes of *Irf1^fl/fl^* and T cell-specific *Irf1*-KO mice were activated by CD3 and CD28 Abs. Cells were harvested at 0, 6, 12, and 24 hours, then subjected to RNA-sequencing. As shown by principal component analysis (PCA), samples group together based on genotype, with clear separation between CD8^+^ T cells from *Irf1^fl/fl^Cd4^Cre^*compared to *Irf1^fl/fl^* (Fig. S4).

Differential expression analysis revealed widespread transcriptional dysregulation in IRF1-deficient CD8⁺ T cells that increased over time following TCR engagement (Fig. 5A). At 6 hours, 243 transcripts were upregulated and 729 transcripts were downregulated in IRF1-deficient CD8⁺ T cells compared to controls. The magnitude of transcriptional changes expanded at 12 hours (476 upregulated, 1,619 downregulated) and further at 24 hours (740 upregulated, 1,967 downregulated), indicating a progressive failure to sustain TCR-driven gene expression programs in the absence of IRF1 (Fig. 5A). Volcano plot analysis highlighted reduced expression of multiple genes associated with TCR signaling, transcriptional activation, and effector differentiation, including Nfatc1, Nr4a1, Nr4a2, Rel, Irf4, and Stat1 (Fig. 5B). Consistent with impaired activation, IRF1-deficient CD8⁺ T cells exhibited diminished induction of immediate-early transcription factors critical for T cell activation.

**Fig. 5:**
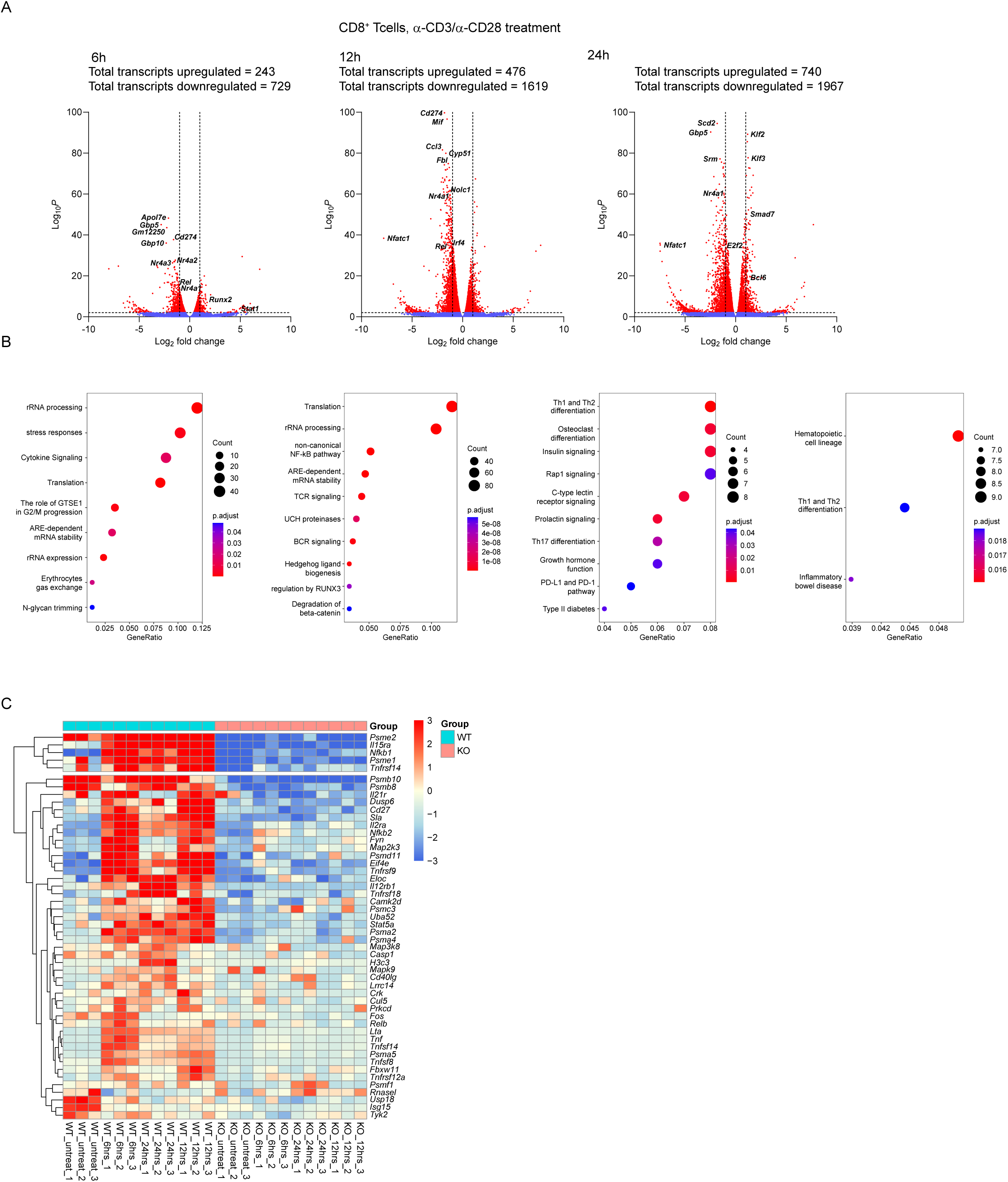
Dysregulation of Nfatc1 expression in T cells from *Irf1^fl/fl^Cd4^Cre^* mice. Purified CD8+ T cells from the spleens of *Irf1^fl/fl^* and *Irf1^fl/fl^Cd4^cre^*mice were activated ex vivo by anti-CD3 and anti-CD28 Abs. Cells were harvested at 0, 6, 12, and 24 hours, then subjected to RNA-sequencing. (A) The volcano plots show the log2FC and −log10(p-value) of select DETs in Naïve CD8+ T cells from mice samples harvested at 6, 12, and 24 hrs in Irf1fl/fl vs Irf1fl/flCd4Cre mice samples. Each datapoint represents a transcript, the red color represents (significant) DETs that have adjusted p-value < 0.05, and log2FC > + 0. The blue color represents DETs that do not match p-value or log2FC cut-off criteria. (B) From left to right the dot plots show the over-represented pathways by genes that are down-regulated and upregulated genes in Irf1fl/fl at 6 hrs harvesting timepoint followed by 12 hrs harvesting timepoint. The overrepresented pathways are annotated in Reactome database for down-regulated genes and in KEGG database for up-regulated genes. (C) The heatmap shows differences in expression of transcription factors that are significantly differentially expressed. The color gradient from blue to red is obtained using z-transformed TPM values, where negative values (blue) identify low expressed genes and vice versa.

Pathway enrichment analyses revealed significant downregulation of pathways related to TCR signaling, NF-κB signaling, cytokine signaling, in addition to translation and ribosomal biogenesis in IRF1-deficient CD8⁺ T cells (Fig. 5B). Notably, TCR signaling and ARE-dependent mRNA stability pathways were among the most significantly enriched pathways affected at later time points, suggesting impaired signal propagation and transcriptional amplification following TCR engagement. Heatmap analysis of selected differentially expressed genes demonstrated consistent suppression of activation-associated transcriptional programs across biological replicates in IRF1-deficient cells (Fig. 5C), further supporting a cell-intrinsic requirement for IRF1 in sustaining TCR-induced gene expression.

### Absence of IRF1 leads to reduced expression of NFAT proteins

We confirmed the findings from RNA-seq data by stimulating naïve CD8+ T cells from wild type and T cell-specific *Irf1*-deficient mice with CD3, CD28 Ab and mouse IL-2, and assessed the protein expression levels of NFATc1 after stimulation. The clear reduction in NFATc1 expression between the two groups was observed as early as 2 hours post-stimulation (Fig. 6A), and the significant difference lasted untill the end of the experiment (6 hours) (Fig. 6A-6B). On the other hand, the expression of Nur77, which is rapidly induced by TCR stimulation as well, was detected and compared as an internal control. As shown in Fig. 6C-D, Nur77 levels increased quickly after the TCR stimulation, and were comparable between the wild type and *Irf1*-KO CD8^+^ T cells. The data indicates that IRF1 deficiency results in reduced NFATc1 protein expression after TCR stimulation leading to compromised CD8^+^ T cell activation.

**Fig. 6:**
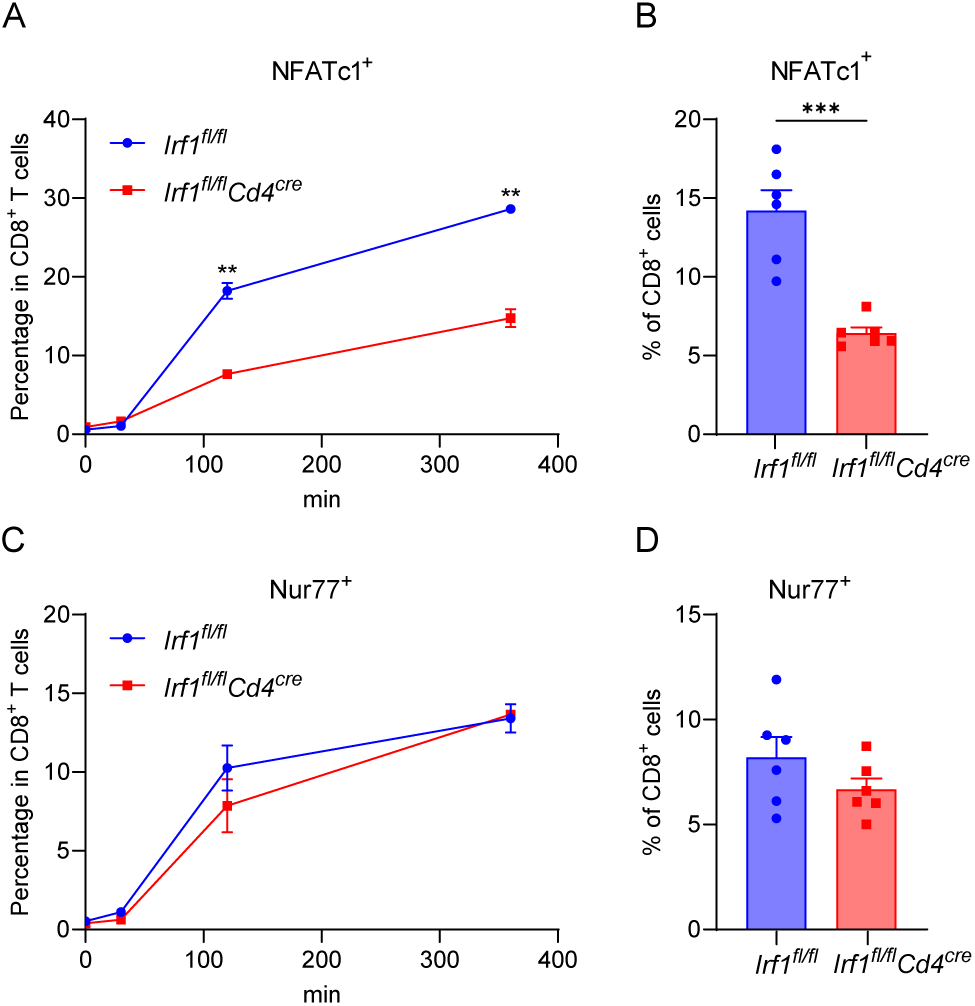
The expression of NFATc1 decreases in *Irf1*-KO CD8^+^ T cells. Naïve CD8+ T cells were isolated from splenocytes of *Irf1^fl/fl^* and *Irf1^fl/fl^Cd4^Cre^*mice using MojoSort mouse CD8 naïve T cell isolation kit, then treated with mouse anti-CD3 Ab, anti-CD28 Ab and IL-2, and cells were harvested and subjected to flow staining at 30 min, 120 min and 360 min (0.5 hr, 2 hrs and 6 hrs). (A) The expression of NFATc1 was measured at different time points after treatment by flow cytometry. (B) Percentage of NFATc1^+^ in CD8+ T cells at 6 hours post-treatment. (C) The expression of Nur77 was measured at different time points after treatment by flow cytometry. (D) Percentage of Nur77^+^ in CD8^+^ T cells at 6 hours post-treatment. The experiments were repeated twice. For each data point, mean and SEM were plotted, and statistical significance was calculated by unpaired t-test, and presented as *** P < 0.001.

### IRF1 expression is correlated with NFATC1 expression in infiltrated T-cells in human melanoma

To confirm our findings in human melanoma, we examined single cell RNAseq data (23) for the localization of T cells within the melanoma tumor microenvironment using UMAP projection and overlaid IRF1 and NFATC1 expression to evaluate their spatial distribution. Both transcripts were enriched within the T-cell compartment and exhibited overlapping regions of high expression, consistent with coordinated regulation in tumor-infiltrating T cells (Fig. 7A-B). To formally quantify this relationship, we calculated per-sample pseudobulk expression values and assessed correlations at the patient level. Across all T cells, IRF1 expression showed a strong positive correlation with NFATC1 (r = 0.81, p = 0.0082) (Fig. 7C). This association was further strengthened when restricting the analysis to CD8⁺ T cells (r = 0.85, p = 0.0035), indicating that the IRF1–NFATC1 axis is particularly prominent within the cytotoxic T-cell compartment (Fig. 7D). In contrast, the correlation within CD4⁺ T cells was weaker and did not reach statistical significance (r = 0.49, p = 0.263), suggesting cell type–specific regulation (Fig. S5A).

**Fig. 7.**
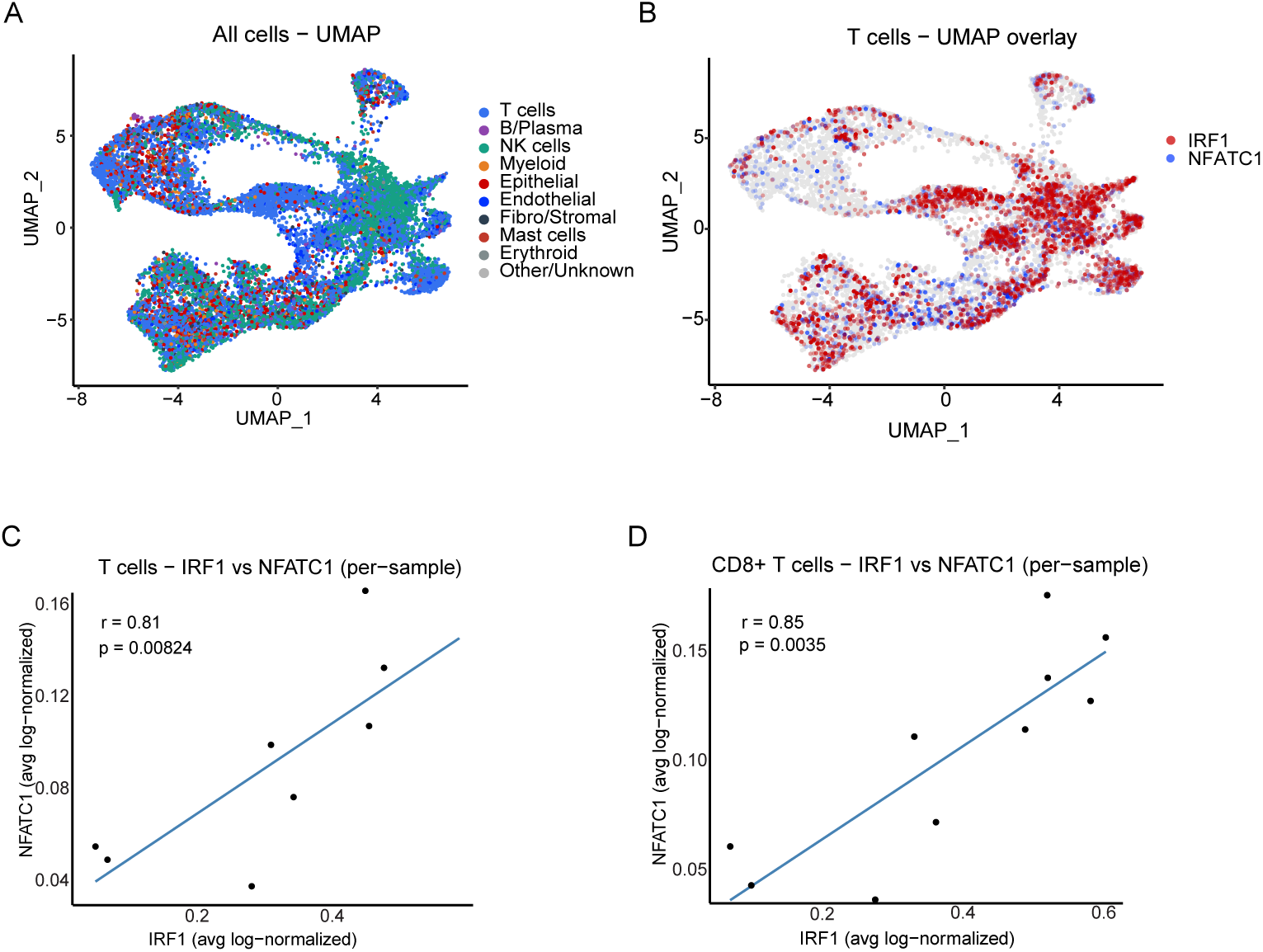
Correlation between IRF1 and NFATC1 expression in single-cell RNA-seq data from human melanoma. (A) Uniform manifold approximation and projection (UMAP) of all cells from a human melanoma single-cell RNA-seq dataset (GEO:GSE14819), colored by major cell types, including T cells, B/plasma cells, NK cells, myeloid cells, epithelial cells, endothelial cells, fibroblast/stromal cells, mast cells, erythroid cells, and other/unknown populations. (B) UMAP displaying overlaid expression of IRF1 (red) and NFATC1 (blue), highlighting regions of co-enrichment within the T-cell compartment. (C) Per-sample pseudobulk analysis across all T cells showing the relationship between sample-level mean (log-normalized) IRF1 and NFATC1 expression assessed by Spearman correlation. Each point represents an individual sample; the line indicates the fitted trend. (D) Per-sample pseudobulk analysis restricted to CD8⁺ T cells showing sample-level mean IRF1 versus NFATC1 expression, assessed by Spearman correlation. Each point represents an individual sample; the line indicates the fitted trend.

To determine whether this relationship reflected a broader NFAT transcriptional program, we extended the analysis to additional NFAT family members. Among these, NFATC1 exhibited the strongest association with IRF1 (ρ = 0.72, p = 0.024), followed by moderate correlations with NFATC3 (ρ = 0.62) and NFATC2 (ρ = 0.58). In contrast, NFATC4 (ρ = 0.17) and NFAT5 (ρ = 0.30) demonstrated minimal association with IRF1, indicating relative specificity of the IRF1–NFATC1 relationship within tumor-infiltrating T cells (Fig. S5B). Taking together these results validated our findings that IRF1 plays a critical role in promoting T cell activation through NFATC1 upregulation in human melanoma.

## DISCUSSION

IFNs and IRFs form a tightly coupled signaling network that exerts profoundly context-dependent effects in cancer and metastasis (6). Canonically, IFNs through JAK–STAT pathways, induce ISGs and IRFs, which together enhance tumor immunogenicity by promoting antigen presentation, cytostasis, and apoptosis, while orchestrating innate and adaptive antitumor immunity in early tumorigenesis (24). However, the same pathways can become tumor-promoting depending on signal strength, duration, and microenvironmental context: chronic or low-grade IFN signaling drives immunoediting, selection of IFN-insensitive clones, upregulation of inhibitory ligands (e.g., PD-L1), and establishment of an immunosuppressive niche that supports metastasis and therapy resistance (25). Thus, IFN–IRF networks act less as linear antitumor pathways and more as adaptive, bidirectional regulators whose impact is dictated by timing, cellular compartment, and tumor type.

IRF1 is a pleiotropic transcription factor that integrates interferon signaling with cell-intrinsic and immune cell–mediated defense programs. IRF1 controls constitutive and inducible expression of host defense genes, linking PRR signaling to cytokine production, antigen presentation, and effector differentiation across innate and adaptive compartments (26). However, as discussed above, recent observations of cell-type and context-specific contributions of IRF1 have necessitated the use of tissue-specific IRF1-deficient mice to pinpoint its function in different cancers. In colorectal cancer models, it functions in both epithelial and myeloid compartments to restrain tumorigenesis by driving PANoptosis, highlighting a tumor-suppressive, immunoregulatory role that is tightly coupled to cell death pathways (27). In viral infection settings, such as MHV68 IRF1 acts as an amplifier of antiviral immunity in T cells (28). Our findings in this context illuminate a molecular mechanism of IRF1’s role in T cell activation. In accordance with previous reports, we found that unlike the germline IRF1-deficient mice, T cell specific loss of IRF1 does not result in severe loss of CD8⁺ T cells (28). However, IRF1 is needed for complete T cell activation. Following TCR stimulation downstream expression of NFATc1 is significantly reduced in the absence of IRF1 resulting in reduced T cell activation and tumor infiltration. Analysis of human melanoma transcriptomic data corroborated this finding establishing a conserved role of IRF1 in NFATc1 expression in T cells. As NFATc1 controls cytotoxicity of CD8⁺ T cells this may be one of the mechanisms underlying the enhanced tumor growth found in *Irf1^fl/fl^CD4^Cre^*mice (29).

The NFAT family of transcription factors are of primary importance during T cell activation and differentiation (30,31). Similar to IRF family transcription factors, NFAT proteins also display significant cell and tissue type-specific expression. Cooperative binding of NFAT and IRF to composite promoters has already been shown to activate various immune cell specific cytokines such as IL-4, IL-10, and IL-12 (32,33). Consistent with its function, NFATc1 expression is known to be autoregulated and also regulated by NFATc2. (34,35). Additionally, NF-κB has been reported to regulate NFATc1 expression during Osteoclast differentiation (36).

Our study suggests the requirement of IRF1 for full expression of NFATc1. Although we do not know the detailed molecular mechanism of this regulation, it is possible that IRF1 and NF-κB may cooperate for complete activation of the NFATc1 promoter, which has been described before in the context of HIV long terminal repeat enhancer (37). Beyond CD8^+^ cells, NFAT proteins have been implicated in the function of CD4^+^ cells, particularly in the suppressive activity of Treg cells (31). However, we did not see significant difference in the Treg population in the tumor microenvironment, which was further supported by our observation that NFATc1 and IRF1 expression did not show significant correlation within CD4⁺ T cells. Collectively, our data uncover a previously unappreciated IRF1–NFATc1 axis in T cells, providing a mechanistic link between interferon signaling and the transcriptional control of T cell activation with functional consequences for antitumor immunity.

## ACKNOWLEDGMENTS

This work was supported in part by AI150214, and AI176333 from NIH, as well as funding from UMGCCC through the Maryland Department of Health’s Cigarette Restitution Fund Program – CH-649-CRF and the National Cancer Institute - Cancer Center Support Grant (CCSG) – P30CA134274.

## AUTHOR CONTRIBUTIONS

LS and SNS designed and conceived the project. LS with help from HB and LK carried out all the experiments. IM and JD carried out the RNAseq data analysis, while EU and ARB contributed in scRNAseq data analysis of human melanoma.

## SUPPLEMENTATRY FIGURE LEGENDS

**Fig. S1:**
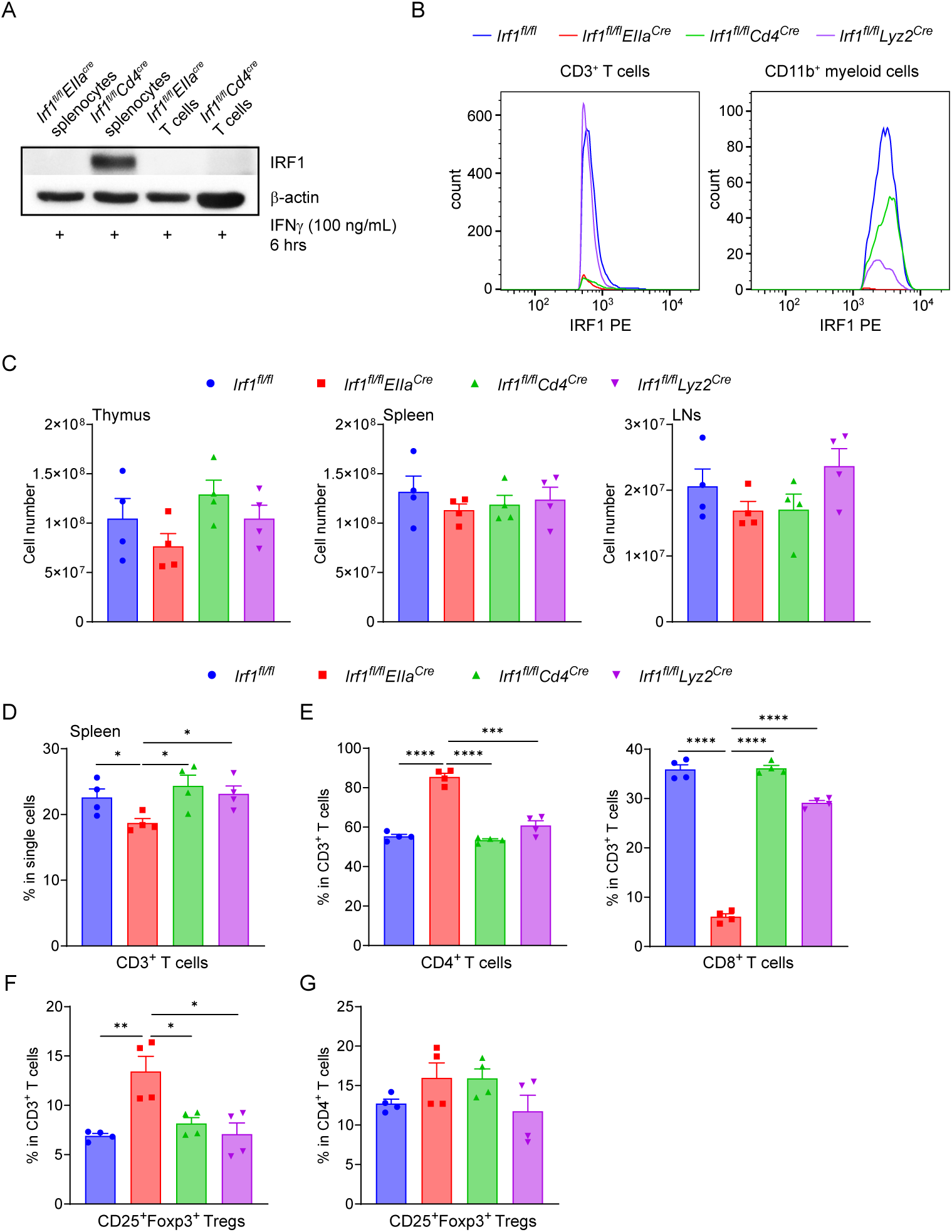
Confirmation of the knock-out of IRF1 in different immune cells and different T cell populations in tissue-specific *Irf1*-KO mice. (A) Splenocytes and isolated T cells from *Irf1^fl/fl^EIIa^Cre^*and *Irf1^fl/fl^Cd4^Cre^* mice were treated with mouse IFNγ, then cell lysates were harvested to confirm the deletion of IRF1 by western blotting. (B) Splenocytes from *Irf1^fl/fl^, Irf1^fl/fl^EIIa^Cre^*, *Irf1^fl/fl^Cd4^Cre^* and *Irf1^fl/fl^Lyz2^Cre^*mice were treated with mouse IFNγ and subjected to flow staining to confirm the removal of IRF1 in myeloid cells and T cells. (C) The total cell number of each thymus, spleen and LNs from different tissue-specific *Irf1*-KO mice. (D) The percentage of CD3^+^ T cells in spleen and (E) the percentage of CD4^+^ T cells and CD8^+^ T cells in CD3^+^ T cells. The percentage of Treg in CD3^+^ T cells (F) and in CD4^+^ T cells (G).

**Fig. S2:**
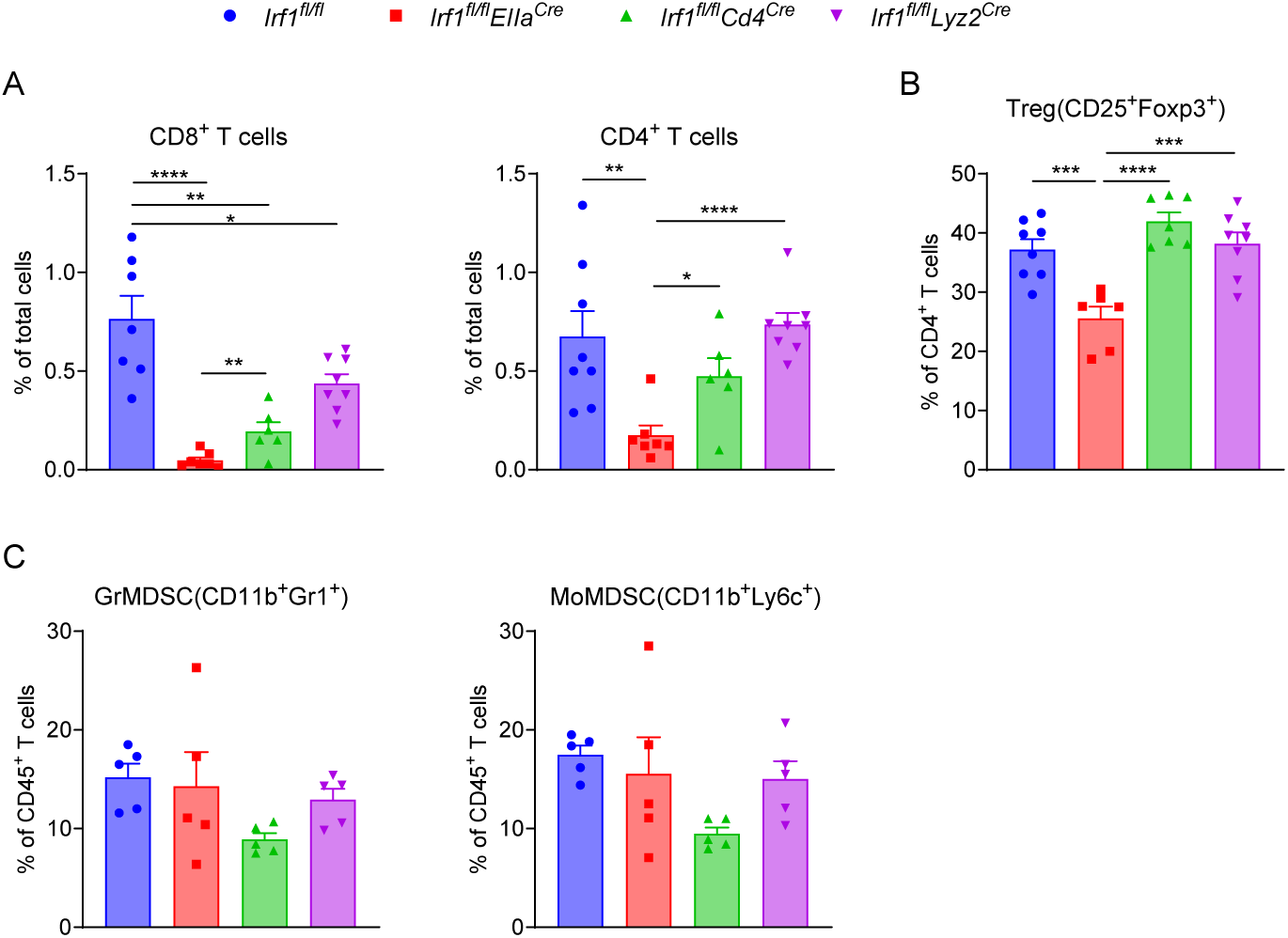
Tumor infiltrating lymphocytes in B16-F10 tumors of different tissue-specific *Irf1*-KO mice. (A) The percentages of CD8^+^ and CD4^+^ T cells in total tumor cells. (B) The percentage of Treg in CD4^+^ T cells. (C) The percentages of GrMDSC and MoMDSC in CD45^+^ cells.

**Fig. S3:**
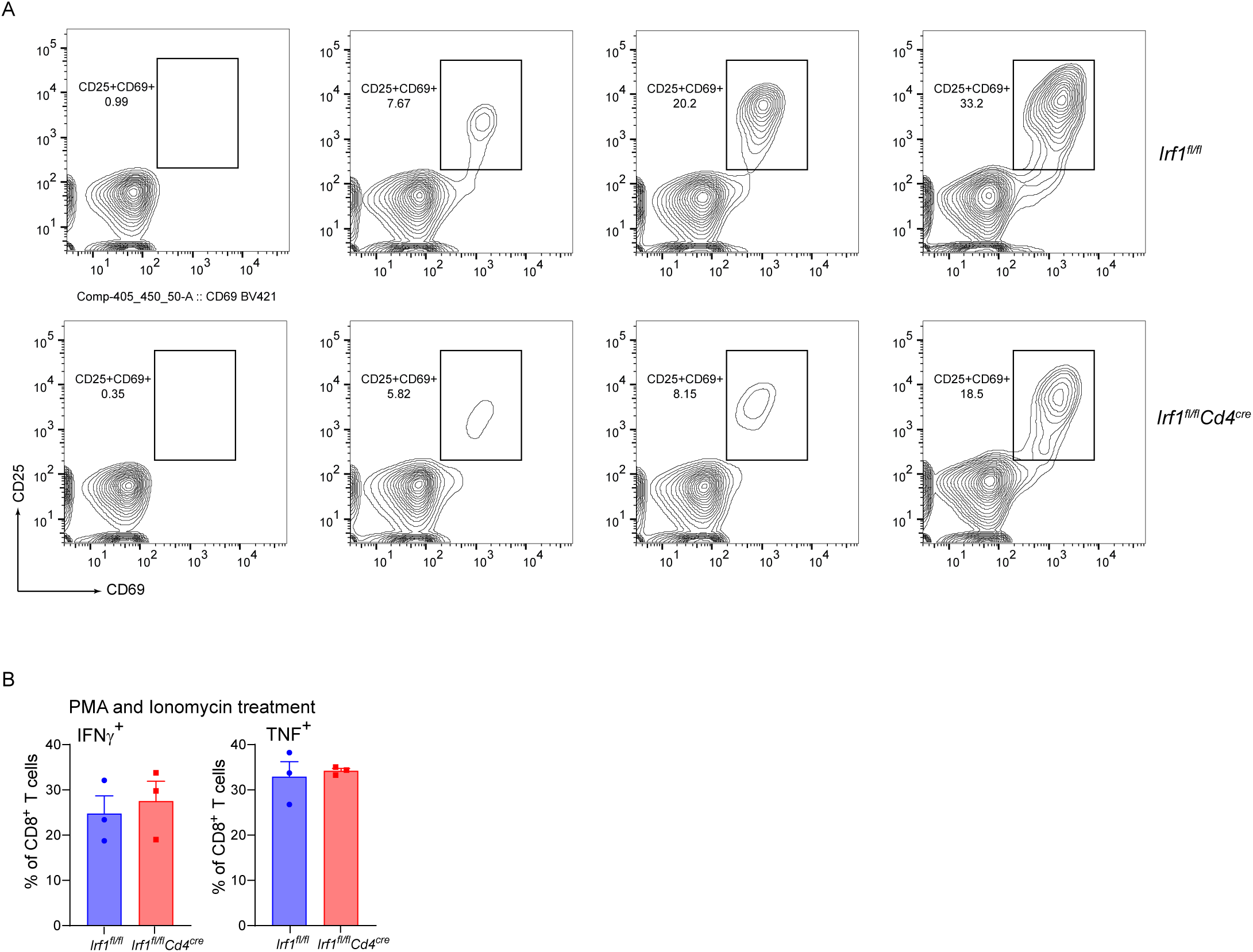
Cytokine productions are not changed in Irf1-KO CD8+ T cells when activated by PMA and ionomycin. Naïve CD8^+^ T cells were isolated from splenocytes of *Irf1^fl/fl^* and *Irf1^fl/fl^Cd4^Cre^*mice and the purity of the isolated CD8^+^ T cell was examined via flow cytometry, then treated with PMA and ionomycin, and cells were harvested and subjected to flow staining (A). (B) Gating plot of CD25^+^ and CD69^+^ in WT and Irf1-KO CD8^+^ T cells at 0 hr, 6 hrs, 12 hrs and 24 hrs after stimulation.

**Fig. S4:**
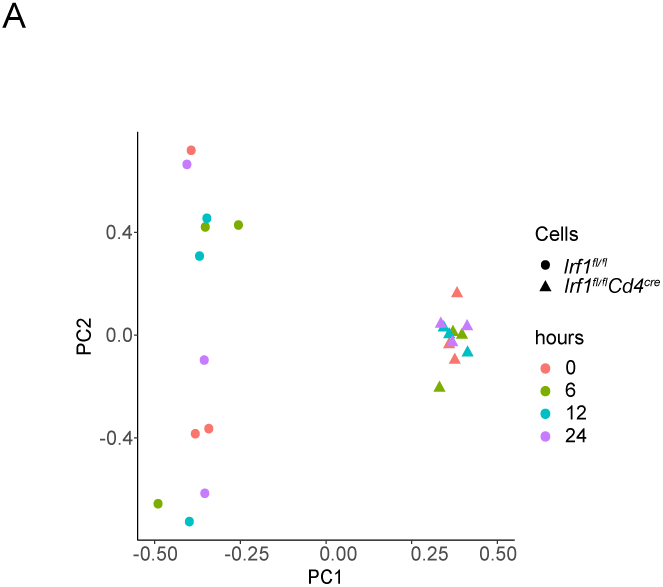
Principal Component analysis of RNAseq data. Principal component analysis (PCA) plot showing the variance between CD8+ T cells from Irf1fl/flCd4Cre when compared to the Irf1fl/fl mice samples harvested at 0, 6, 12, and 24 hrs after stimulation. The shape of the dots represents the groups and the colors represent the harvesting time.

**Fig. S5.**
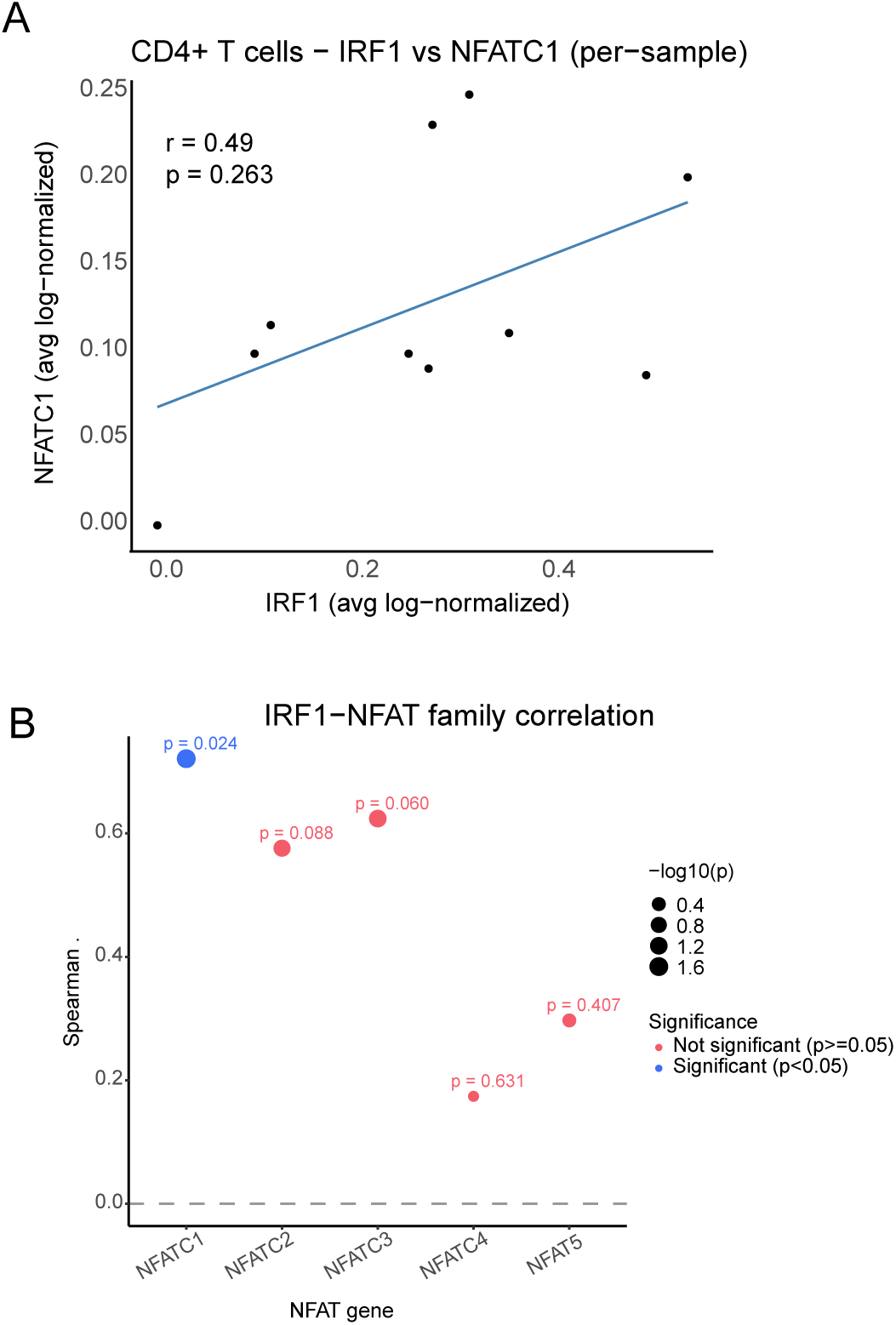
IRF1–NFAT relationships and NFATC1-associated T-cell state shifts in a human melanoma single-cell RNA-seq dataset (GSE148190). (A) Per-sample pseudobulk scatter plot illustrating the relationship between IRF1 and NFATC1 expression within CD4⁺ T cells. Each point represents an individual sample; the fitted line denotes the linear trend. The Spearman correlation coefficient (*r*) and corresponding *p* value are shown. (B) Dot plot summarizing Spearman correlations between IRF1 and each NFAT family member (NFATC1, NFATC2, NFATC3, NFATC4, NFAT5) using per-sample pseudobulk averages computed from tumor-infiltrating T cells in the melanoma single-cell RNA-seq dataset. Point color denotes nominal significance (blue, *p* < 0.05; red, *p* ≥ 0.05), and point size scales with−log10(*p*); nominal *p* values are indicated adjacent to each point.

## REFERENCES

1. Taniguchi T, Ogasawara K, Takaoka A, Tanaka N. IRF family of transcription factors as regulators of host defense. Annu Rev Immunol. 2001;19:623–55.

2. Tanaka N, Taniguchi T. The interferon regulatory factors and oncogenesis. Seminars in Cancer Biology. 2000;10:73–81.

3. Negishi H, Taniguchi T, Yanai H. The Interferon (IFN) Class of Cytokines and the IFN Regulatory Factor (IRF) Transcription Factor Family. Cold Spring Harb Perspect Biol. 2018;10:a028423.

4. Willman CL, Sever CE, Pallavicini MG, Harada H, Tanaka N, Slovak ML, et al. Deletion of IRF-1, mapping to chromosome 5q31.1, in human leukemia and preleukemic myelodysplasia. Science. 1993;259:968–71.

5. Alsamman K, El-Masry OS. Interferon regulatory factor 1 inactivation in human cancer. Bioscience Reports. Portland Press Ltd; 2018;38:BSR20171672.

6. Tamura T, Yanai H, Savitsky D, Taniguchi T. The IRF Family Transcription Factors in Immunity and Oncogenesis. Annual Review of Immunology. Annual Reviews; 2008;26:535–84.

7. Van Der Weyden L, Arends MJ, Campbell AD, Bald T, Wardle-Jones H, Griggs N, et al. Genome-wide in vivo screen identifies novel host regulators of metastatic colonization. Nature. 2017;541:233–6.

8. Karki R, Sharma BR, Lee E, Banoth B, Malireddi RKS, Samir P, et al. Interferon regulatory factor 1 regulates PANoptosis to prevent colorectal cancer. JCI Insight [Internet]. 2020;5. Available from: http://www.ncbi.nlm.nih.gov/pubmed/32554929

9. Shao L, Hou W, Scharping NE, Vendetti FP, Srivastava R, Roy CN, et al. IRF1 Inhibits Antitumor Immunity through the Upregulation of PD-L1 in the Tumor Cell. Cancer Immunology Research. 2019;7:1258–66.

10. Shao L, Srivastava R, Delgoffe GM, Thorne SH, Sarkar SN. An IRF2-Expressing Oncolytic Virus Changes the Susceptibility of Tumor Cells to Antitumor T Cells and Promotes Tumor Clearance. Cancer Immunol Res. 2024;12:779–90.

11. Penninger JM, Sirard C, Mittrücker HW, Chidgey A, Kozieradzki I, Nghiem M, et al. The interferon regulatory transcription factor IRF-1 controls positive and negative selection of CD8+ thymocytes. Immunity. 1997;7:243–54.

12. Lohoff M, Mak TW. Roles of interferon-regulatory factors in T-helper-cell differentiation. Nature Reviews Immunology. Nature Publishing Group; 2005;5:125–35.

13. Bray NL, Pimentel H, Melsted P, Pachter L. Near-optimal probabilistic RNA-seq quantification. Nat Biotechnol. Nature Publishing Group; 2016;34:525–7.

14. Pimentel H, Bray NL, Puente S, Melsted P, Pachter L. Differential analysis of RNA-seq incorporating quantification uncertainty. Nat Methods. Nature Publishing Group; 2017;14:687–90.

15. Subramanian A, Tamayo P, Mootha VK, Mukherjee S, Ebert BL, Gillette MA, et al. Gene set enrichment analysis: A knowledge-based approach for interpreting genome-wide expression profiles. Proceedings of the National Academy of Sciences. Proceedings of the National Academy of Sciences; 2005;102:15545–50.

16. Wu T, Hu E, Xu S, Chen M, Guo P, Dai Z, et al. clusterProfiler 4.0: A universal enrichment tool for interpreting omics data. The Innovation. 2021;2:100141.

17. Kanehisa M, Goto S. KEGG: kyoto encyclopedia of genes and genomes. Nucleic Acids Res. 2000;28:27–30.

18. Gillespie M, Jassal B, Stephan R, Milacic M, Rothfels K, Senff-Ribeiro A, et al. The reactome pathway knowledgebase 2022. Nucleic Acids Res. 2022;50:D687–92.

19. Hao Y, Hao S, Andersen-Nissen E, Mauck WM, Zheng S, Butler A, et al. Integrated analysis of multimodal single-cell data. Cell. 2021;184:3573–3587.e29.

20. McInnes L, Healy J, Saul N, Großberger L. UMAP: Uniform Manifold Approximation and Projection. Journal of Open Source Software. 2018;3:861.

21. Lee JH, Lee BH, Jeong S, Joh CS-Y, Nam HJ, Choi HS, et al. Single-cell RNA sequencing identifies distinct transcriptomic signatures between PMA/ionomycin- and αCD3/αCD28-activated primary human T cells. Genomics Inform. 2023;21:e18.

22. Courtney AH, Lo W-L, Weiss A. TCR Signaling: Mechanisms of Initiation and Propagation. Trends Biochem Sci. 2018;43:108–23.

23. Mahuron KM, Moreau JM, Glasgow JE, Boda DP, Pauli ML, Gouirand V, et al. Layilin augments integrin activation to promote antitumor immunity. J Exp Med. 2020;217:e20192080.

24. von Locquenghien M, Rozalén C, Celià-Terrassa T. Interferons in cancer immunoediting: sculpting metastasis and immunotherapy response. J Clin Invest. 2021;131:e143296, 143296.

25. Ruiz-Iglesias A, Guilbaud E, Galluzzi L, Mañes S. Context-dependent impact of type I interferon signaling in cancer. Mol Cancer. 2025;24:275.

26. Feng H, Zhang Y-B, Gui J-F, Lemon SM, Yamane D. Interferon regulatory factor 1 (IRF1) and anti-pathogen innate immune responses. Blumenthal A, editor. PLOS Pathogens. PLoS Pathog; 2021;17:e1009220.

27. Karki R, Sharma BR, Lee E, Banoth B, Malireddi RKS, Samir P, et al. Interferon regulatory factor 1 regulates PANoptosis to prevent colorectal cancer. JCI insight [Internet]. JCI Insight; 2020;5. Available from: http://www.ncbi.nlm.nih.gov/pubmed/32554929

28. Jondle CN, Johnson KE, Mboko WP, Tarakanova VL. T Cell-Intrinsic Interferon Regulatory Factor 1 Expression Suppresses Differentiation of CD4+ T Cell Populations That Support Chronic Gammaherpesvirus Infection. J Virol. 2021;95:e0072621.

29. Klein-Hessling S, Muhammad K, Klein M, Pusch T, Rudolf R, Flöter J, et al. NFATc1 controls the cytotoxicity of CD8+ T cells. Nat Commun. 2017;8:511.

30. Macian F. NFAT proteins: key regulators of T-cell development and function. Nat Rev Immunol. Nature Publishing Group; 2005;5:472–84.

31. Müller M, Rao A. NFAT, immunity and cancer: a transcription factor comes of age. Nat Rev Immunol. 2010;10:645–56.

32. Farrow MA, Kim E-Y, Wolinsky SM, Sheehy AM. NFAT and IRF Proteins Regulate Transcription of the Anti-HIV Gene, *APOBEC3G*\*. Journal of Biological Chemistry. 2011;286:2567–77.

33. Rengarajan J, Mowen KA, McBride KD, Smith ED, Singh H, Glimcher LH. Interferon Regulatory Factor 4 (IRF4) Interacts with NFATc2 to Modulate Interleukin 4 Gene Expression. J Exp Med. 2002;195:1003–12.

34. Zhou B, Cron RQ, Wu B, Genin A, Wang Z, Liu S, et al. Regulation of the murine Nfatc1 gene by NFATc2. J Biol Chem. 2002;277:10704–11.

35. Chuvpilo S, Jankevics E, Tyrsin D, Akimzhanov A, Moroz D, Jha MK, et al. Autoregulation of NFATc1/A expression facilitates effector T cells to escape from rapid apoptosis. Immunity. 2002;16:881–95.

36. Kim JH, Kim N. Regulation of NFATc1 in Osteoclast Differentiation. J Bone Metab. 2014;21:233–41.

37. Sgarbanti M, Remoli AL, Marsili G, Ridolfi B, Borsetti A, Perrotti E, et al. IRF-1 is required for full NF-kappaB transcriptional activity at the human immunodeficiency virus type 1 long terminal repeat enhancer. J Virol. 2008;82:3632–41.

